# Identifying target sites for placental therapeutics through the comparison of normal term pregnancies and intrauterine growth restricted proteomes

**DOI:** 10.1101/2021.12.15.472796

**Authors:** A. Bowman, H. Brockway, H. Jones

## Abstract

A variety of pathologies, including intrauterine growth restriction (IUGR), have been linked to placental insufficiencies as important causal factors, however, little has been done to develop therapeutics that may improve placental development, structure and function. The placenta offers the opportunity to manipulate the in-utero environment without directly intervening with the fetus, accessible from the maternal circulation, a vital but temporary organ, the placenta is no longer required after birth. Developing therapeutics must involve multiple aspects of targeting and safety to ensure no off-target impact on the pregnant person or fetus as well as enhance efficiency of delivery. In addition to our studies on nanoparticle delivery to the placenta [1] we are developing targeting strategies to allow peripheral delivery via the maternal circulation. In this study we have performed the isolation of the microvillous membrane from human placental syncytiotrophoblast (the targeting cell) and via proteomic analysis identified potential targeting moieties, we have then compared this to publicly available data from pregnancies complicated by fetal growth restriction to ensure that we do not identify targets which would be reduced in FGR, resulting in less efficient targeting.

**Highlights:**

- Proteomic analysis detected key genes, proteins, and pathways present in the syncytiotrophoblast membrane of normal placentas.
- Specific membrane proteins identified show promise for future characterization of placental pathologies, such as IUGR.
- Proteomes of normal and diseased placentas require further study to better understand the etiology of certain conditions.

## Introduction

Intrauterine growth restriction (IUGR) is defined as a condition where fetal birth weight is less than the tenth percentile for any given gestational age, indicating the fetus is not reaching its recommended growth targets during gestation. Growth restriction leads to increased health risks both during infancy and long-term [2]. Interestingly, IUGR can also occur with preeclampsia (PE), an adverse pregnancy outcome characterized by maternal hypertension and proteinuria. In comparison to preeclampsia’s effects on pregnant mothers (upwards of 18% of maternal death), IUGR is associated with high mortality rates in infants with 10% of perinatal mortality linked to the condition [3]. IUGR is typically diagnosed via Doppler ultrasound [5]. Despite advances in obstetric care, most cases of IUGR result in iatrogenic preterm birth (induced birth less than 37 weeks gestational age) due to fetal distress or, in cases associated with PE, maternal distress [4].

Some studies point to placental insufficiencies as a contributor to both preeclampsia and IUGR, however, further evaluation is crucial to determine the precise etiology of the placental insufficiency [2, 5]. Recently, systems biology approaches and the resulting analyses of “omics” data have advanced placental research to explore pathologies such as IUGR and PE at a fundamental level. Preeclampsia and IUGR “omics” suggest altered gene expression and/or genetic mutation could possibly contribute to the condition [6]. The purpose of our innovative study was to compare and contrast the placental membrane proteomics of normal term pregnancies to establish placental sufficiency under optimal conditions, then to identify publicly available IUGR and/ PE+IUGR omics including proteomics and transcriptomes for comparison. This would allow us to identify key proteins and pathways that have essential roles in placental sufficiency processes. Our work is novel in that we are directly observing proteins found in the syncytiotrophoblast membrane of the human placenta. We hypothesized that placentas from IUGR and PE cases will have distinct protein and gene expression in identified membrane targets from that found in normal term placentas.

While transcriptome expression in placentas has been utilized as a way to compare and contrast pathologies with normal physiological processes, there is a current lack of placental proteomic data from both normal and adverse pregnancy outcomes. This represents a significant gap in the knowledge surrounding normal placental physiology and the molecular mechanisms that may be at play surrounding maternal-fetal conditions [7].

## Materials and Methods

### Isolation of syncytial MVM vesicles

De-identified human placentas (n=3) from term deliveries (38-42 weeks gestational age) with no reported complications were obtained from the University of Cincinnati Medical Center (UCMC). Isolation of placental syncytial trophoblast membranes were performed according to previously optimized methods [8].

### Proteomics

50 μg of isolated vesicles was run in three separate wells on a 4-12% NT NuPage gel using MOPS running buffer. After the gel was ran the bands were excised and reduced using dithiothreitol (DTT), alkylated with lodoacetamide and digested with trypsin. The resultant peptides were recovered, dried in a speed vacuum, and resuspended in 125 μL of 0.1% Formic acid (FA). From here 2 μg of each peptide was analyzed using nanoscale liquid chromatography (nanoLC-MS) tandem mass spectrometry. For each sample, a list of duplicate peptides (99% confidence) was generated, and proteins were punitively identified via comparison with the Uniprot protein *Homo sapiens* database [9] using the Protein Pilot program (Sciex). The protein % coverage inferred the presence of protein in the membranes which was then averaged to determine a threshold for further analyses. Proteins with a >30% coverage were run back through Uniprot for identification and further annotated with Biomart [10]. To identify common proteins across all three membrane samples we compared the individual proteins lists and utilized proteins that were represented across all three samples for further analysis.

### Characterization of protein interactions

The individual protein list formed from the normal term microvillous membranes were submitted to STRING [11], a protein-protein interaction network, to identify proven protein interactions revealed by experimental data, databases, co-expression, co-occurrence, and gene fusion. The network of proteins was then clustered through a kmeans clustering of 3 under the highest interaction confidence (0.900), which is a setting within the String program creating predicted protein networks based on known protein interactions with confident research evidence.

### Biological pathway analysis

Proteins identified with >20% coverage were submitted for PANTHER [12] overrepresentation tests under *Homo sapiens* reference corrected by Fisher’s Exact test with Bonferroni, looking at Reactome and Panther pathways with respective fold enrichments and adjusted P-values (p>0.05 Fishers Test with Bonferroni Correction for multiple comparisons). This 20% was used as a cutoff coverage for various analyses performed to provide accurate results for larger populations of proteins present.

Next a cross-reference of pathways between our omics data and the public FGR datasets was performed. The FGR dataset was ran through PANTHER [12] under the same conditions as our omics for both Reactome and Panther pathways. A cross-reference using Microsoft Excel was performed to compare the pathways present within the two datasets.

### Comparison to IUGR omics

The individual protein list formed from the normal term placentas was uploaded to QIAGEN’s Ingenuity Pathway Analysis [21] program where a comparative analysis was performed against differentially expressed proteins from a published study assessing the proteomes of both fetal growth restricted and normal pregnancies [22]. The resulting list of overlayed molecules were then complied into a network which was formatted to show its molecular interactions on a subcellular level (Figure 2). A cross-reference of the two sets showed no similarities between the proteins in either dataset.

## Results

### Common protein expression

From 291 individual proteins identified (S1 Table) in the placental membranes from normal term pregnancies, 171 of these had an average percent coverage >20%. The identified proteins were matched with their respective estimated expression to observe relationships between transcription and translation. Proteins with the highest estimated expressions were analyzed alongside their corresponding percent coverage (Table 1).

**Table 1.** Highly expressed proteins from placentas with >20% coverage.

| **Protein name** | **Average coverage (%)** | **Expression estimation data (RPKM)** |
| --- | --- | --- |
| CSH2 | 58.373334 | 9176.96579 |
| EF1A3 | 35.5699996 | 1319.49231 |
| CP19A | 41.5500005 | 1188.46578 |
| 3BHS1 | 49.3300021 | 508.015056 |
| HBB | 91.6133344 | 383.722839 |
| PPB1 | 59.6866687 | 380.258927 |
| GTR1 | 21.7466667 | 338.223669 |
| S100P | 60.3533328 | 332.588116 |
| EZRI | 41.4100001 | 273.770204 |
| TGM2 | 42.7466671 | 243.184123 |
| ANXA1 | 75.5300005 | 225.077402 |
| CAN6 | 42.0166671 | 215.930713 |
| ACTB | 74.4866669 | 213.561799 |
| HBA | 80.5166662 | 191.948339 |
| NIBL1 | 23.1000001 | 191.464606 |
| CALX | 25.6766667 | 185.078137 |
| HBG2 | 92.5199986 | 184.05798 |
| TFR1 | 40.7433331 | 182.368248 |
| AHNK | 21.163334 | 166.844181 |
| CLIC4 | 51.910001 | 161.697758 |

### Distinctive protein interactions

Our complete proteomic dataset contains several identified proteins of interest that are related to placental function and IUGR including: cell division control protein 42 (CDC42), ras-related protein Rab-5C (RAB5C), complement glycoprotein encoded by CD59 (CD59), and spectrin beta protein (SPTB). We took a closer look into percent coverage and gene expression for each of these proteins (Table 2). Although the proteins had >20% coverage, the gene expression for RAB5C and SPTB was quite low suggesting these are not actively being transcribed in contrast to expression levels for CD59 and CDC42.

**Table 2.** Proteins associated with IUGR identified in our dataset.

| Proteins | Average coverage (%) | Expression estimate (RPKM) |
| --- | --- | --- |
| CD59 | 26.56 | 113.02 |
| CDC42 | 42.41 | 61.02 |
| RAB5C | 32.56 | 7.78 |
| SPTB | 24.60 | 1.03 |

Furthermore, this protein set was unique in the sense that they also demonstrated a high number of protein-protein interactions within the membrane of the normal term placentas itself, which was identified by running all of the proteins through the STRING database (Figure 1).

**Figure 1.**
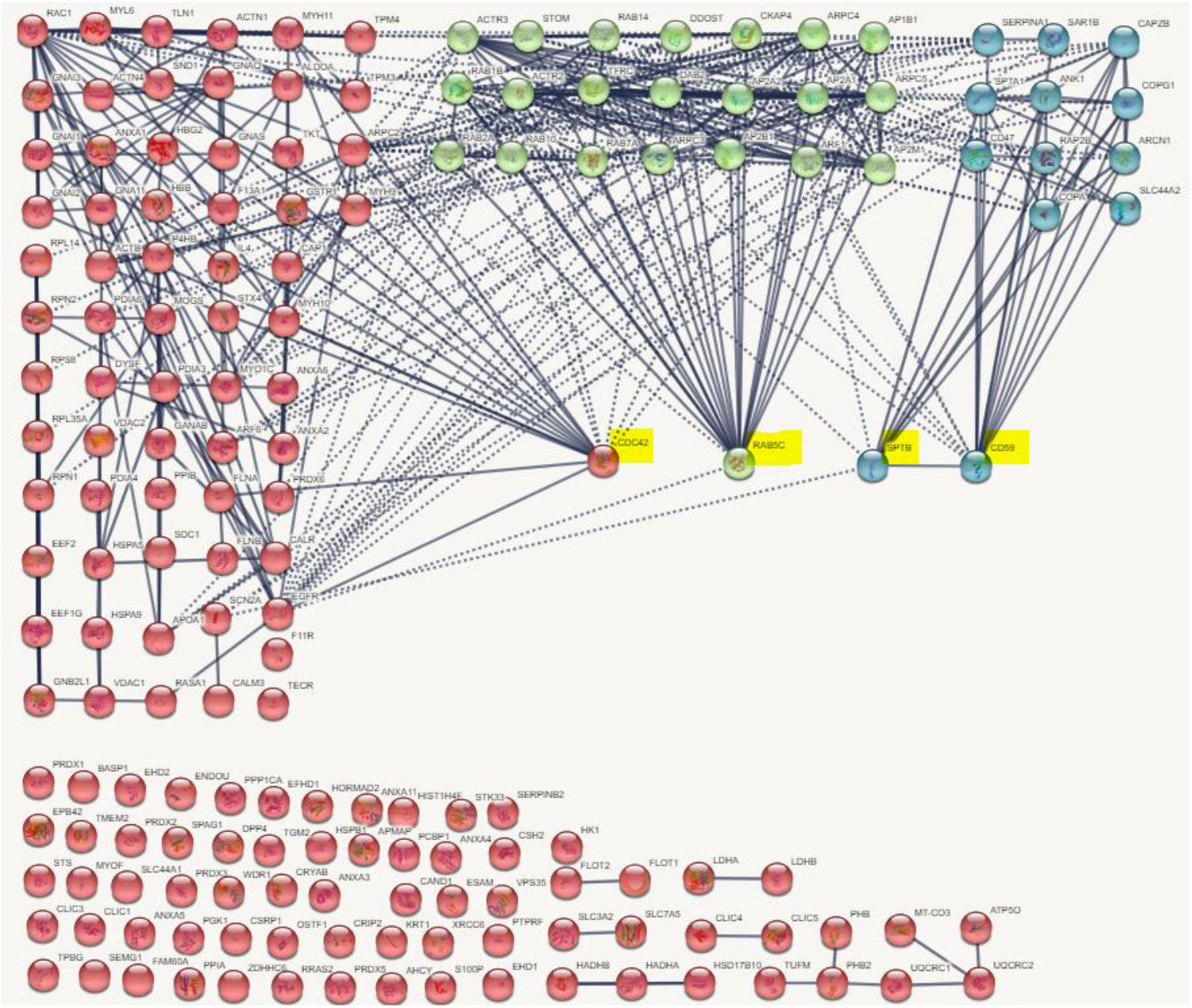
Protein interactions: STRING data demonstrates the proteins of interest have a high number of protein-protein interactions, suggesting an essential network of protein networks that may have a role in placental function and sufficiency. The line thickness is representative of the strength of data supporting the interaction between the present proteins, dashed lines being indicative of less data support.

Interestingly, our selected proteins also interact with key receptors, calreticulin (CALR) and epidermal growth factor receptor (EGFR) (Figure 1). CALR and EGFR are membrane receptor proteins with a significant number of interactions amongst the other membrane proteins. However, other receptor proteins such as, F11-receptor (F11R) and trans-2,3-enoyl-CoA reductase (TECR) show no proven interaction with other proteins present within the placental membrane proteome (Figure 1).

### Expression of enriched biological pathways

PANTHER pathway analyses identified several molecular pathways with higher-than-expected fold enrichments including glycolysis, cytoskeleton regulation, endogenous cannabinoid signaling, and PI3 kinase (Table 3). Many of these significant pathways play vital roles in cell growth, cell signaling, hormone mediation and disease prevention, which are all important for a healthy pregnancy.

**Table 3.** Identified Panther pathways from proteins with >20% coverage.

| **PANTHER Pathways** | **Total proteins in the pathway** | **Proteins from our dataset** | **Fold Enrichment** | **Adj P value\*** |
| --- | --- | --- | --- | --- |
| Cytoskeletal regulation by Rho GTPase | 83 | 12 | 17.53 | 2.86E-09 |
| Inflammation mediated by chemokine and cytokine signaling pathway | 255 | 15 | 7.13 | 1.03E-06 |
| PI3 kinase pathway | 52 | 8 | 18.65 | 4.68E-06 |
| Glycolysis | 19 | 6 | 38.28 | 6.59E-06 |
| Huntington disease | 143 | 11 | 9.33 | 8.98E-06 |
| Gonadotropin-releasing hormone receptor pathway | 232 | 12 | 6.27 | 1.29E-04 |
| Integrin signaling pathway | 191 | 10 | 6.35 | 9.82E-04 |
| Dopamine receptor mediated signaling pathway | 57 | 6 | 12.76 | 0.002 |
| Opioid prodynorphin pathway | 33 | 5 | 18.37 | 0.002 |
| Opioid proopiomelanocortin pathway | 34 | 5 | 17.83 | 0.003 |
| Opioid proenkephalin pathway | 34 | 5 | 17.83 | 0.003 |
| Enkephalin release | 35 | 5 | 17.32 | 0.003 |
| 5HT1 type receptor mediated signaling pathway | 46 | 5 | 13.18 | 0.009 |
| Metabotropic glutamate receptor group II pathway | 48 | 5 | 12.63 | 0.01 |
| Endogenous cannabinoid signaling | 24 | 4 | 20.2 | 0.01 |
| EGF receptor signaling pathway | 136 | 7 | 6.24 | 0.03 |
| Muscarinic acetylcholine receptor 2 and 4 signaling pathway | 61 | 5 | 9.94 | 0.03 |
| Corticotrophin releasing factor receptor signaling pathway | 32 | 4 | 15.15 | 0.03 |
| Parkinson disease | 98 | 6 | 7.42 | 0.03 |
| GABA-B receptor II signaling | 36 | 4 | 13.47 | 0.05 |

<sup>a</sup> Fishers test with Bonferroni Correction for multiple comparisons.

The PANTHER overrepresentation test was used to identify present reactome pathways in the normal term placentas with >20% coverage, discovering numerous pathways related to RHO GTPases, Interleukin signaling, cellular degranulation, and Insulin-like Growth Factor (IGF) transport. Of the 97 reactome pathways identified (S2 Table) significant recurrences in the aforementioned pathways were spotlighted (Table 4) due to their active roles in cellular proliferation, cellular development, and fetal immune response.

**Table 4.** Observed reactome pathways from proteins with >20% coverage.

| Reactome Pathways | Total proteins in the pathway | Proteins from our dataset | Fold Enrichment | Adj P value* |
| --- | --- | --- | --- | --- |
| RHO GTPases activate PAKs (R-HSA-5627123) | 21 | 5 | 28.86 | 0.004 |
| RHO GTPases Activate WASPs and WAVes (R-HSA-5663213) | 35 | 7 | 24.25 | 9.52E-05 |
| RHO GTPases activate IQGAPs (R-HSA-5626467) | 31 | 6 | 23.46 | 0.001 |
| RHO GTPases activate PKNs (R-HSA-5625740) | 63 | 9 | 17.32 | 1.51E-05 |
| RHO GTPase Effectors (R-HSA-195258) | 287 | 20 | 8.45 | 1.97E-09 |
| Signaling by Rho GTPases (R-HSA-194315) | 417 | 20 | 5.81 | 1.12E-06 |
| Gene and protein expression by JAK-STAT signaling after Interleukin-12 stimulation (R-HSA-8950505) | 37 | 8 | 26.21 | 5.87E-06 |
| Interleukin-12 signaling (R-HSA-9020591) | 46 | 9 | 23.72 | 1.27E-06 |
| Interleukin-12 family signaling (R-HSA-447115) | 56 | 10 | 21.65 | 3.04E-07 |
| RAB geranylgeranylation (R-HSA-8873719) | 65 | 7 | 13.06 | 0.004 |
| Signaling by Interleukins (R-HSA-449147) | 447 | 18 | 4.88 | 1.13E-04 |
| Innate Immune System (R-HSA-168249) | 1105 | 39 | 4.28 | 3.48E-11 |
| Cytokine Signaling in Immune system (R-HSA-1280215) | 823 | 25 | 3.68 | 5.78E-05 |
| Platelet degranulation (R-HSA-114608) | 127 | 15 | 14.32 | 1.53E-09 |
| Response to elevated platelet cytosolic Ca<sup>2+</sup> (R-HSA-76005) | 132 | 15 | 13.78 | 2.55E-09 |
| GTP hydrolysis and joining of the 60S ribosomal subunit (R-HSA-72706) | 112 | 11 | 11.91 | 1.15E-05 |
| Platelet activation, signaling and aggregation (R-HSA-76002) | 259 | 25 | 11.7 | 1.79E-15 |
| Neutrophil degranulation (R-HSA-6798695) | 478 | 27 | 6.85 | 1.9E-11 |
| Regulation of Insulin-like Growth Factor (IGF) transport and uptake by Insulin-like Growth Factor Binding Proteins (IGFBPs) (R-HSA-381426) | 124 | 8 | 7.82 | 0.027 |

<sup>a</sup> Fishers test with Bonferroni Correction for multiple comparisons.

The cross-reference of Reactome and Panther pathways from the FGR data [22] and our study’s omics showed a few pathways that are present in both datasets. The Reactome pathways present in both sets include Regulation of Insulin-like Growth Factor (IGF) transport, Neutrophil degranulation, and Innate immune system. The Panther pathway that the two sets shared was the Integrin signaling pathway.

Comparison of the present proteomes from both the normal term placentas in our study and the publicly available fetal growth restricted data [22] allowed for the creation of an overlayed network of the following molecules: LPCAT3, RPL18A, ARF6, DPP4, EIF4A1, EPB42, HBB, HBG2, HSP90B1, IQGAP2, ITGAV, IgG1, IgG1, P4HB, PRDX2, RAP2B, RASA1, RPL18, S100A10, SERPINA1, SERPINB2, SLC4A1, SPTA1, SPTB. This network was connected and formatted to demonstrate the interactions within this network on a subcellular molecular level (Figure 2). Although many of the molecules within this subcellular network are self-regulating some, such as SPTB, Hbb-b1, and SERPINA1 show interactions among others within the overlayed network.

**Figure 2.**
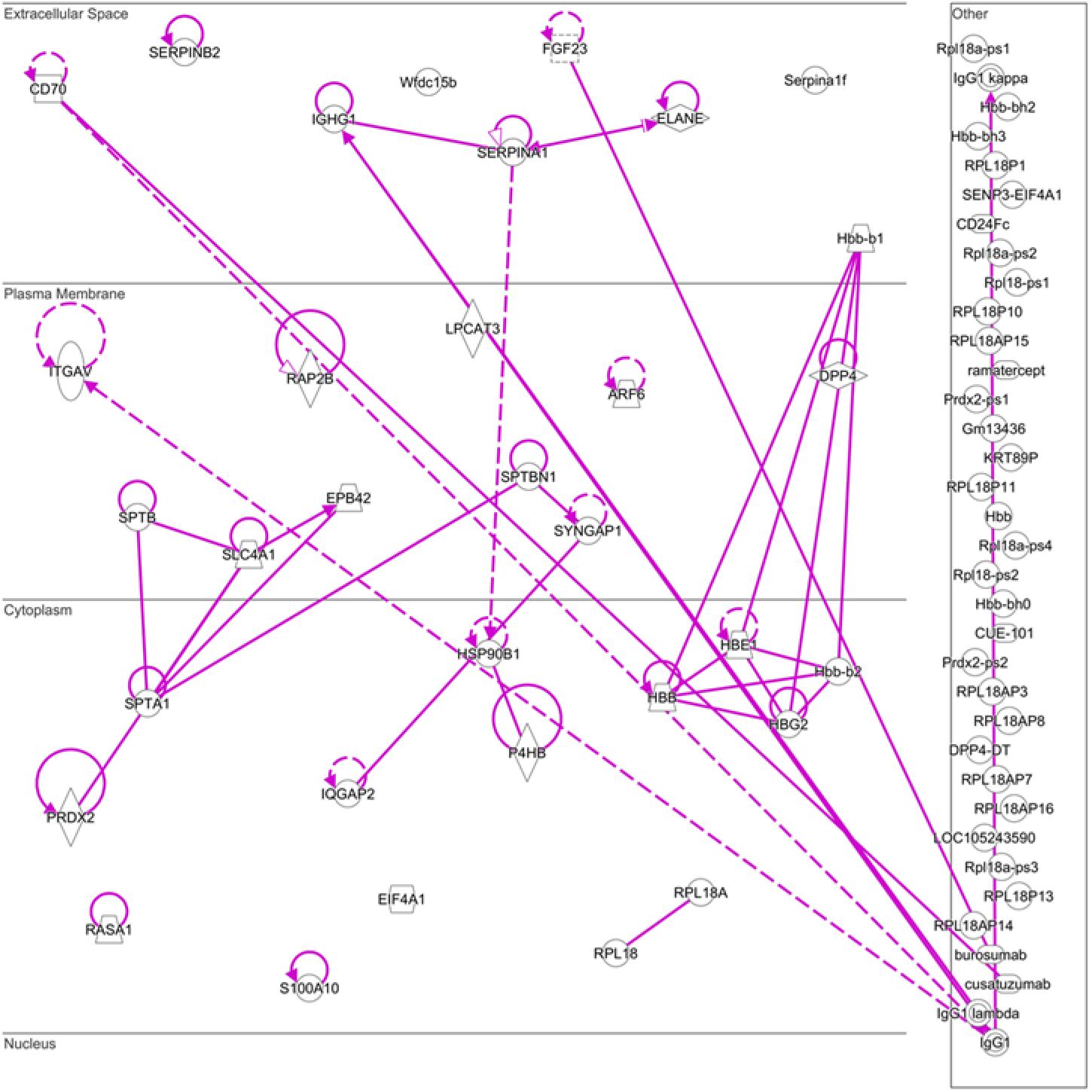
Subcellular interactions of overlayed proteins: Ingenuity Pathway Analysis comparison between our normal term protein list and that of publicly available proteome data from fetal growth restricted research shown in the form of subcellular molecular interactions. These interactions show promise in identifying potentially important networks within fetal growth restricted placentas.

## Discussion

Within our dataset we were able to identify 171 membrane proteins with >20% coverage, several of which have been previously associated with adverse pregnancy outcomes including IUGR. Our research (Table 2) supports previous data showing normal term pregnancies have high expression of CD59 and abnormal function of this protein is linked to placental pathologies such as IUGR and PE [13] CD59 plays an important role in preventing C5b-9 membrane attack complex (C5b-9MAC), whose presence in pregnancy is associated with cases of IUGR and PE [13]. We found high average percent coverage and expression of protein CDC42 in our normal term placenta sample (Table 2) while previous studies report finding decreased expression and activity in cases of IUGR [14]. CDC42 is a Rho-GTPase family protein that aids in developing cellular structure and regulation of the actin cytoskeleton [13] which is important to note as the syncytiotrophoblast membrane is comprised of microvilli that are necessary for appropriate membrane function including transport and signaling [18]. The Rho-GTPase family falls under the larger characterization of RAS superfamily proteins that have downstream effects signaling genes involved in cell growth, differentiation, and vesicular trafficking [14]. One of the RAS superfamily proteins observed in our samples include RAB5C. RAB5C plays a role in vesicular trafficking which is crucial to understanding IUGR as reports show a decrease in microvilli in pregnancies with this pathology [15]. Our normal sample data also had low expression levels of protein SPTB which is involved in the formation of the cytoskeletal structure of erythrocyte plasma membranes. It is unclear if this is a part of the syncytial membrane or if it is representing the presence of fetal and maternal blood in the sample. Although this is interesting concept, additional data is necessary before any conclusions can be drawn.

We were interested in exploring the interactions of these particular proteins, as several were known to be active in signaling pathways, specifically, CD59 and CDC42 due to their high expression levels (Table 2) and their previous association to IUGR and other placental pathologies. Our data showed the numerous interactions shown between CD59 and CDC42 with other proteins present in a normal placental membrane (Figure 1). Both had interactions with other proteins responsible for cytoskeletal development and molecule transport. CDC42 showed confident interactions with proteins Guanine nucleotide-binding protein subunit alpha i1(GNAI1) and Guanine nucleotide-binding protein subunit alpha i2 (GNAI2) involving transmembrane signal transduction [11], this supports our findings of several molecular signaling pathways in normal pregnancies including regulation of cytoskeleton (Table 3). Interestingly, CD59 had multiple protein interactions with proteins involved with cytoskeletal mechanisms affecting the erythrocytic membrane and inhibition of thrombin. Proteins such as Ankyrin 1erythrocytic (ANK1), spectrin alpha erythrocytic 1 (SPTA1), and Serpin family E member 1(SERPIN1) point to CD59’s ability to impact the red blood cell membrane and impact rates of coagulation [11]. The high expression rate of CD59 we found in normal term placentas may suggest the importance of platelet degranulation and platelet signaling pathways (Table 4). It is known that hematological changes do occur during pregnancy to accommodate the physiological needs of both mother and fetus [19].

Our, analyses revealed interactions with several key receptors especially calreticulin (CALR) and epidermal growth factor receptor (EGFR) which are linked to similar biological pathways as our other highlighted proteins. CALR is a functioning chaperone that is thought to be localized to the endoplasmic reticulum (ER) and promotes homeostasis [11]. ER stress has been linked to several placental pathologies including PE [20]. Furthermore, CALR is essential to placental function via its role in Ca2+ regulation [20]. EGFR interacts with CDC42 and RAB5C, through its functions in several signaling cascades [11]. Previous studies conducted on EGFR indicate that mutations in this receptor have been observed in cases of IUGR and other pregnancy complications [16] and due to the significantly present enrichment fold of this receptor pathway in normal placentas (Table 3) it should be observed when looking at IUGR proteomes/transcriptomes.

Many of these receptors are responsible for cellular metabolic processes, cellular proliferation, cellular signaling, and response to hormones. The binding affinities for each of the specific receptor proteins present are essential for the appropriate molecular pathway activity to take place. The pathway analyses suggested roles for these proteins in cytoskeletal development, inflammatory regulation, and cellular signaling, all essential to placental function. Cytoskeletal regulation by Rho GTPase pathways, as well as inflammation mediated by chemokine and cytokine signaling pathways observed (Table 3) correlate to the many RHO GTPase pathways and immune system/ Interleukin pathways identified (Table 4). As we are observing similar pathway association in both analyses, we feel confident that these proteins are essential to placental function and require further study.

Our proteomics data when compared to publicly available FGR omics showed no similar proteins shared, however, the Ingenuity Pathway Analysis [21] identified molecules involved in both datasets and mapped them on a subcellular level (Figure 2). This data is useful for potential drug therapeutic targets since these molecules play are present in both normal and growth restricted placentas. These molecules may be contributing to the pathway similarities we identified through our cross-reference of pathway data. Both normal and growth restricted omics have shown pathways with regulation of IGF, neutrophil degranulation, innate immune system, and integrin signaling which points to further exploration of underlying molecules involved in these processes. Given that lowered expression of Insulin-like growth factor (IGF) is associated with IUGR [17], it is interesting that the fold expression of the IGF regulation pathway, as well as the other similar pathways had higher fold enrichments than those from the normal omics data. Additional studies of these specific pathways and the molecules involved in their processes could prove beneficial for comparison of normal and growth restricted placentas and finding targets for drug therapeutics in cases of IUGR.

With a recognized lack of proteomes without complications available for comparison our data contributes support to standardization of proteomes available in placentas from normal term pregnancies. Our recognition of highlighted genes, proteins, and pathways show indication of distinct markers that may play a role in pathologies leading to adverse pregnancy outcomes such as IUGR and PE. Future works are crucial to comparing the placental membrane proteomes and transcriptomic data of normal term pregnancies and those with IUGR and PE, as this would promote earlier clinical diagnoses and better treatment plans for pregnant mothers.

## Acknowledgments

This research was made possible by grants from the University of Cincinnati’s Ronald E. McNair program, the University of Cincinnati’s University Research Council, and the Eunice Kennedy Shriver NICHDR01HD090657.

Mass spectrometry data was collected and analyzed in the UC Proteomics Laboratory under the direction of KD Greis, PhD. Funding for the Sciex 5600 nanoLC-MS/MS system was obtained in part through an NIH shared instrumentation grant (S10 RR027015-01).

## Supporting information

**S1 Table. Complete identification of proteins in normal term isolated Syncytial MVM vesicles.**

(DOCX)

**S2 Table. Complete identification of significant reactome pathways with high fold expression.**

(DOCX)

